# Spatiotemporal Integration in Plant Tropisms

**DOI:** 10.1101/506147

**Authors:** Yasmine Meroz, Renaud Bastien, L. Mahadevan

## Abstract

Tropisms, growth-driven responses to environmental stimuli, cause plant organs to respond in space and time and reorient themselves. Classical experiments from nearly a century ago reveal that plant shoots respond to the integrated history of light and gravity stimuli rather than just responding instantaneously. We introduce a temporally non-local response function for the dynamics of shoot growth formulated as an integro-differential equation whose solution allows us to qualitatively reproduce experimental observations associated with intermittent and unsteady stimuli. Furthermore, an analytic solution for the case of a pulse light stimulus expresses the response function as function of experimentally tractable variables for the phototropic response of *Arabidopsis* hypocotyls. All together, our model enables us to predict tropic responses to time-varying stimuli, manifested in temporal integration phenomena, and sets the stage for the incorporation of additional effects such as multiple stimuli, gravitational sagging etc.

---

Plant tropisms are the growth-driven responses of a plant organ which reorients itself in the direction of an environmental stimulus such as light, termed phototropism, or gravity, termed gravitropism. Tropisms driven by a directional stimulus lead to the asymmetric redistribution of a growth hormone such as auxin [1-5] which then directs growth. For example, in Fig. 1a we show snapshots of the negatively gravitropic response of a wheat seedling placed horizontally at time *t* = 0, where the seedling shoot detects the direction of gravity and grows to oppose it. This response is dynamical, and one might suspect that if the stimulus is changed intermittently, the spatiotemporal response itself will be complex. In the simple experiment described above, gravity acts continuously on the shoot with a constant magnitude. Thus it is not possible to distinguish between a response that integrates the stimulus over time and one that acts instantaneously. However experimental observations of gravitropism and phototropism dating back more than a century have shown that plants respond to time varying stimuli in a way that suggests that they do integrate the stimuli in time. For example, different combinations of stimuli that are intermittent in time [6-10] or which have reciprocal ratios of intensity and duration [11-18], so that the time-integrated stimulus is constant, lead to the same response, as shown in the insets in Figs. 2 and 3. Explanations of shoot phototropism assume that this follows from photobiology [19]. However, the fact that this phenomenon has also been observed in the context of gravitropism suggests that one must look for a common signal transduction pathway, naturally implicating the polar transport of the growth hormone auxin that is critical in mediating tissue growth, which is indeed driven by either gravity or light. Furthermore, these observations of responses to time-varying stimuli also naturally suggest that the plant retains a memory of the stimulus. To quantify this at a minimal level, we turn to linear response theory to characterize the relation between the responsive geometry of growth and the exciting stimulus.

**FIG. 1:**
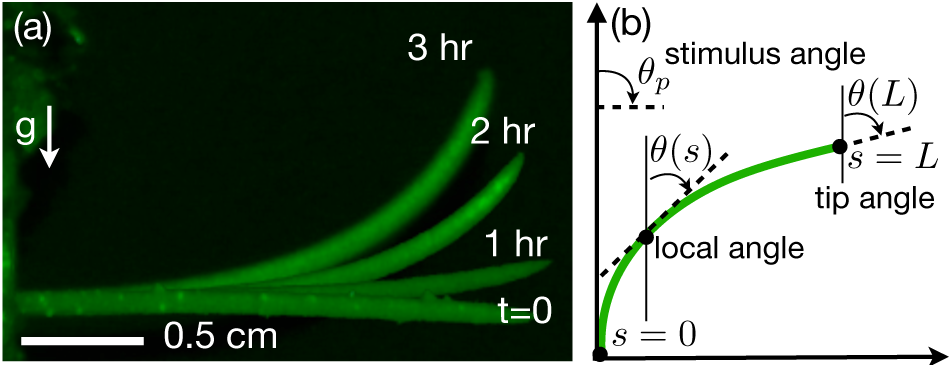
(a) Time course of a gravitropic response of a single wheat seedling placed horizonatally (perpendicular to the direction of gravity) at *t* = 0, and measured at 1 hour intervals. (b) Mathematical definitions [20]. The angle *θ*(*s,t*) at point *s* along the organ at time *t* is defined from the vertical. The parameter *s* runs along the organ from *s* = 0 at the base, to *s* = *L* at the tip. The tip angle *θ*(*L,t*) is the quantity which is conventionally measured in experiments.

**FIG. 2:**
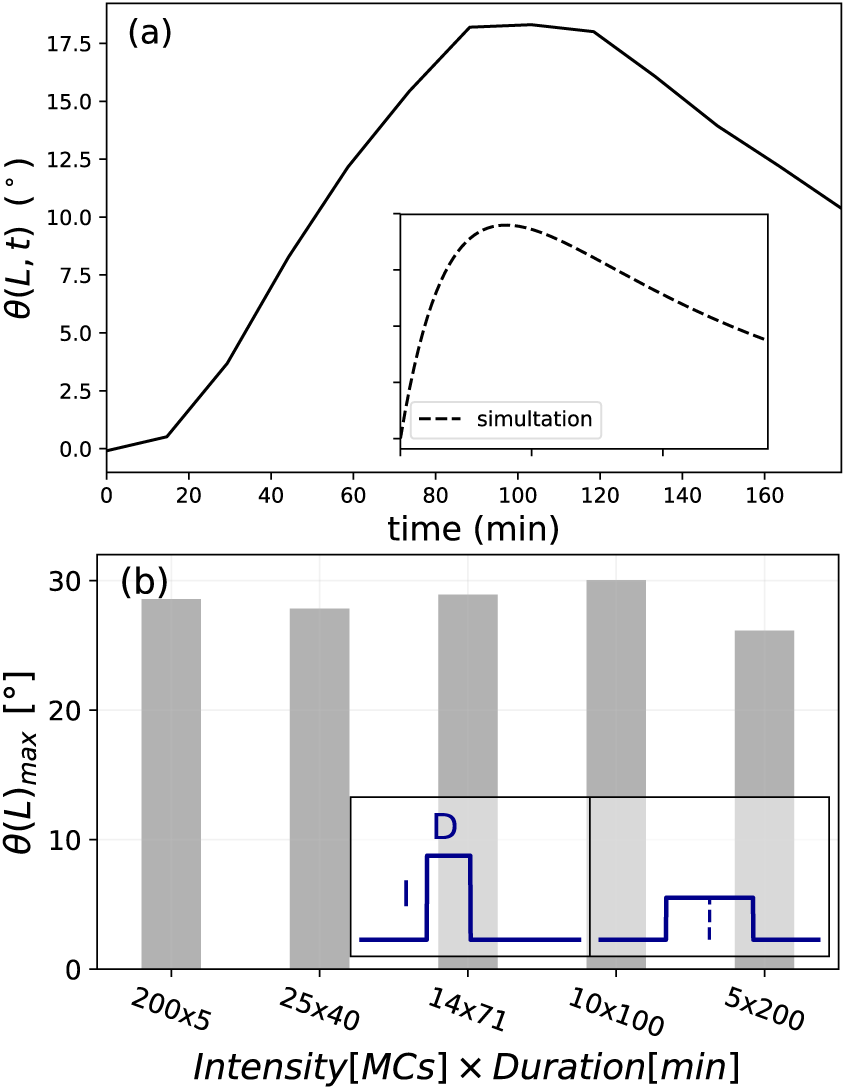
(a) Experimental observations made by Orbovic & Poff [26] who recorded the time course of the tip angle *θ*(*L,t*) of *Arabidopsis Thaliana* seedlings as a response to a 0.9 s pulse of blue light, observed hours after the pulse is gone. The inset shows the simulated response using our response theory approach described in Eq. 2. (b) Reciprocity experiment on the phototropic response of *Avena* coleoptiles, adapted from Briggs (1960) [12]. Inset shows lighting protocols; One unilateral pulse of light *vs* another pulse with half the intensity and double the duration, i.e. identical total dose. The main figure shows the average maximal angle of the tip measured for different reciprocal ratios of intensity (ranging between 5-200 MCs) and duration of exposure time (ranging between 200-5 s), yielding the same total dose of 1000*MC* x *s* ∼ 1.46W/m^2^.

**FIG. 3:**
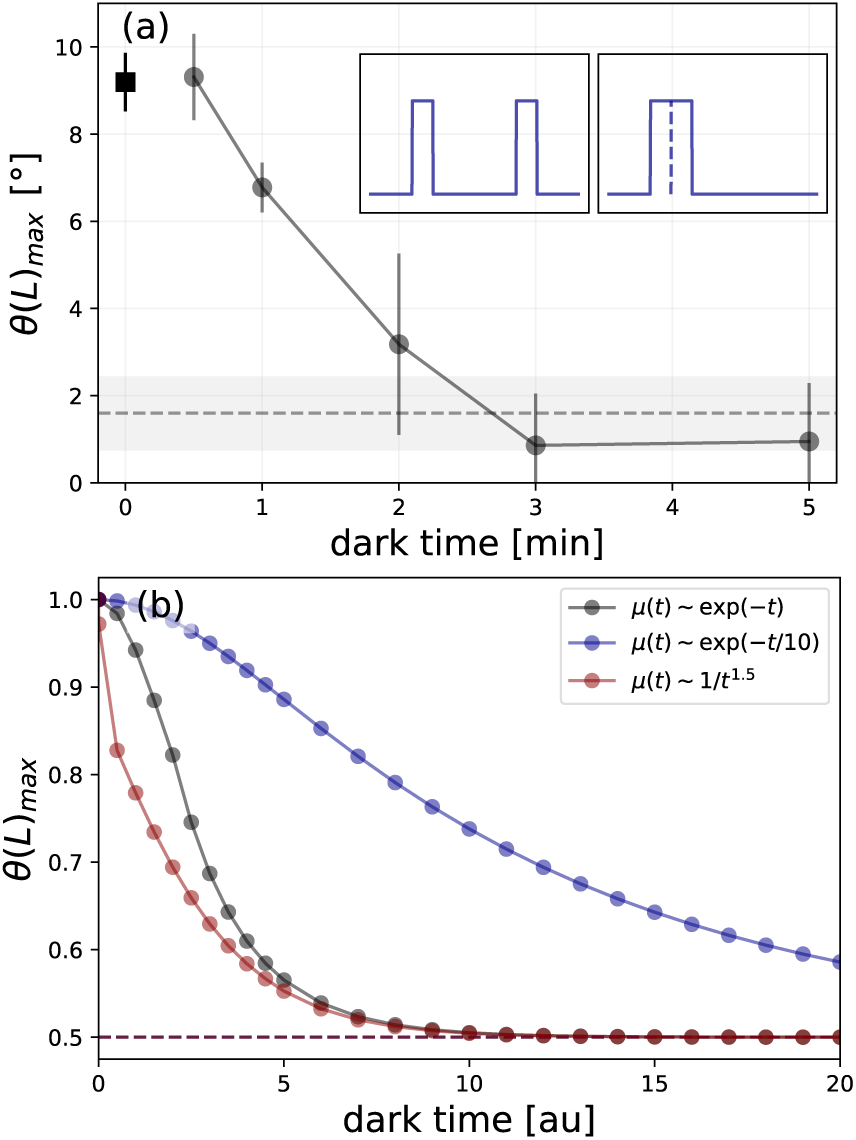
(a) Summation experiment on the phototropic response of *Vaucheria geminata,* adapted from Kataoka (1979) [10]. Inset shows lighting protocols; two pulses of light separated by dark time, versus a single continuous pulse with double the duration. Circles are the average maximal tip angle (vertical bars indicate SE) measured for the response to two light pulses, each 10s and 6*Wm*^-2^, separated by a dark interval of various durations (x-axis). The square symbol is the response to a continuous pulse of 20s (zero lag). The vertical dashed line is the average response to a single pulse of 10s, where the gray bar indicates the SE. (b) Equivalent simulations of summation experiment, using different response functions; two exponentials *μ*(*t*) ∼ exp(−*t/τ*) with ***τ*** = 1 (black) and ***T*** = 10 (blue), and a power-law *μ*(*t*) ∼ 1/*t*^1.5^ (red). Maximal tip angles are normalized to allow comparison, so that the maximal angle for zero lag is always 1, and the control response to a single pulse is 0.5 (dashed line).

The description of input-output relations of a signal transducer characterize the output *y*(*t*) as the weighted sum of the input signal *x*(*t*) convolved by a response function *μ*(*t*), so that we may write 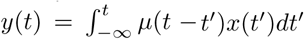. This approach has been used in a variety of problems concerning temporal responses of organisms to external stimuli, including bacterial chemotaxis [21, 22], cellular chemotaxis [23], and the light induced growth response in Phycomyces [24]. Extending this framework to the tropisms seen in plant shoots requires coupling the non-local temporal response to the growth-driven dynamical changes in the shape of the whole plant organ, leading to a spatio-temporal framework that is qualitatively different from previous purely temporal theories, as we will see.

We start from a recent framework describing the kinematics of tropic responses that combines internal proprioception and external phototropism or gravitropism [20, 25] to explain the growth kinematics of shoots subject to uniform and constant stimuli. The shape of a slender shoot growing in a single plane, Fig. 1b, can be described in terms of the local angle *θ*(s, *t*) of the tangent to the shoot from the vertical as a function of the arc-length along the centerline *s* at time t. The shoot actively grows only within its growth zone, of length *L*_gz_ which is smaller than the length of the entire organ *L*. Within the growth zone, *L > s > L − L*_gz_, observations suggest [20] that the rate of change of the curvature in the growth zone is proportional to a weighted sum of the stimulus term associated with the environment, and the proprioceptive term that penalizes deviations from a straight shoot. Then the kinematics of shoot growth for *L > s > L − L*_gz_ follow the equation

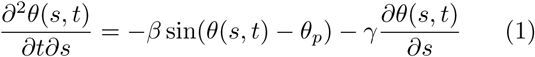

Here *θ*_*p*_ is the angle of the stimulus (light or gravity) relative to the vertical, 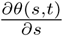 is the local curvature, and the parameters *β* and *γ* are the sensitivities to gravity (or light) and proprioception. The ratio *B* = *βL*_gz_*/γ* is a dimensionless bending response parameter which determines the relative importance of proprioception and (gravi/photo) tropism, with *B* ∈ [0.9 − 9.3], displaying broad intraspecific and interspecific variability [20]. Outside of the growth zone, *s < L − L*_gz_, the shoot does not respond so that *θ*_*st*_ = 0 here. We note that although the model assumes that the response to the stimulus is a weighted sum of the proprioceptive, gravitropic and phototropic stimulus, it is linear and instantaneous, i.e. it cannot account for the experimental observations of temporal integration discussed earlier. To allow for this we introduce a convolution of the external stimulus term with a response function *μ*(t), leading to a modified form of the dynamic law for shoot reorientation:

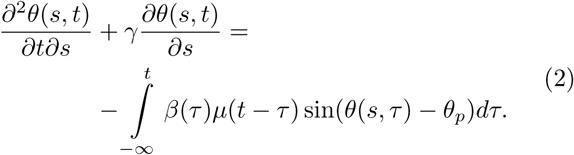

The convolution with the response function *μ*(t) represents the memory associated with the response to a history of stimuli. As one might expect, the experimental observations of reciprocity and summation of stimuli are valid only within some time window, suggesting that the response function should decay with an equivalent characteristic time scale, and needs to be determined experimentally. We note that when *μ*(t) = *δ*(*t*), we recover the original model Eq. 1 corresponding to an instantaneous response.

To see the utility of this modified law, we first turn to the results of an experiment showing the response of *Arabidopsis Thaliana* to a pulse of light, shown in Fig.2(a). We see that the response is observed hours after the pulse is gone, and cannot be described by the instantaneous model Eq. 1 [20]. However, our modified response law in Eq. 2 naturally yields a response occurring long after a stimulus has been switched off, as seen in the simulations in the inset of Fig. 2(a), for a particular choice of the kernel *μ*(t) = *e*^*-–t/10*^ and initial conditions corresponding to a straight seedling, i.e. *∂θ/∂s*|_*t*_*=*_0_ = 0, an initial angle from the vertical *θ*(s, 0) = *θ*_0_ =0, and boundary conditions corresponding to a clamped base, i.e. *θ*(0,t) = 0, and a free end *∂θ/∂s*|_*s=L*_ = 0 while the unilateral lighting stimulus acts at *θ*_*p*_ = π/2. To solve Eq. 2 with these conditions, we use the Verlet integration method [27], with the following parameters: time step of *dt* = 0.005, length of the shoot *L* = 1.0, number of bins dividing the shoot length *N* = 100, so that the spacial element is *ds* = *L/N*, and the gravi/photo-ceptive and proprioceptive sensitivities were taken to be *β* = 5.0 and *γ* = 0.5, at the high end of B values, for numerical efficiency.

Having shown that the integrated response function allows us to capture the response of the growing shoot to a pulse of light, we now turn to try and explain reciprocity experiments. Fig. 2b reproduces reciprocity experiments [12] associated with the phototropic response of *Avena* coleoptiles. The coleoptiles where exposed to identical total dose of unilateral light, but with reciprocal ratios of intensity and exposure time as detailed in the caption of Fig. 2b. The response, measured in terms of the maximal angle of the tip *θ*(*L*)_*max*_, remains identical, showing that the coleoptiles respond to the integrated history of stimuli within this time range. To explain these observations, we use our model as a basis for numerical simulations, emulating the described reciprocity experiments, with the same initial and boundary conditions as for the pulsed light experiment.

Table 1 shows results for simulations employing an exponential response function of the form *μ*(*t*) ∼ exp(−*t*/200), in units of the simulation time step. As expected, these simulations yield similar responses for stimuli with identical doses but reciprocal ratios of intensity and duration, as long as the duration (the longest here being 50) is shorter than the natural decay built into the exponential memory kernel, i.e. ***τ*** = 200.

**TABLE I:**
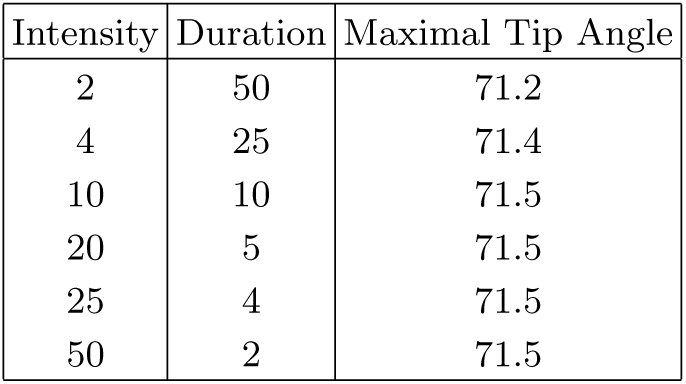
Simulations of reciprocity experiments equivalent to Fig. 2b, employing an exponential response function of the form *μ*(*t*) ∼ exp(−*t*/200), in units of the simulation time step.

This leads us to naturally investigate the integration timescale, and its dependence on the decay of the response function, using a summation experiment in phototropism [10]. In Fig. 3a, we show the maximal angle of the tip *θ*(*L*)_max_, of a coenocytic alga *Vaucheria geminata* to light pulses of 10 seconds each, as a function of dark interval of various durations. We note that for lag times between 30s and ∼ 3m the response decays, going from the response for the sum of both pulses, to that for a single pulse. This decay suggests a smoothly decaying response function, as opposed to a step function, with a timescale of about 2 minutes.

Fig. 3b shows results for simulations employing response functions with different kernels, e.g. an exponential one *μ*(t) ∼ exp(−*t/τ*) with ***τ*** =1,10, and a power-law *μ*(t) ∼ 1/*t*^1.5^. In order to allow a comparison maxi-mal tip angles are normalized so that the maximal angle for zero lag is always unity, and the control response to a single pulse is 0.5. We see that as the intervening dark time increases, the response decays relative to that for a single pulse. Furthermore, our choice of different kernel response functions lead to a different decay in the integrated response, clearly indicating its importance for the correct prediction of tropic responses to dynamic stimuli.

In order to extract the form of the kernel response function from the kinematics of tropic responses, we solve Eq. 2 for the case of a pulse stimulus, by substituting *β*(*t*) = *β*_0_*δ*(*t*_0_ = 0) in the linearized limit. This eliminates the convolution, and together with the initial condition *θ*(*s,t* = 0) = 0 leads to:

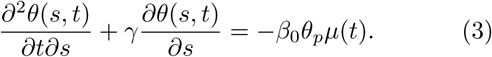

Integrating over *s*, and recalling the boundary condition *θ*(*s* = 0,*t*) = 0, leads to :

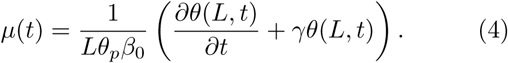

This thus provides an experimentally tractable relation allowing us to extract the kernel response function from the experimentally observed dynamics of the angle at the tip as a response to a pulse stimulus.

Using the measured response of seedlings to a pulse of light [26] shown in Fig. 2a, we now apply Eq. 4 to determine bounds on the kernel response function, using bounds on the time *T*_*c*_ ∈ [80,100] minutes to provide bounds on *γ* ∈ [0.01, 0.0125]. Substituting the measured *θ*(*L,t*), the calculated 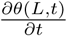 and the two bounds for 7 into Eq. 4, we get two bounds for the functional form, up to a prefactor, of the response function *μ*(t), shown in Fig. 4.

**FIG. 4:**
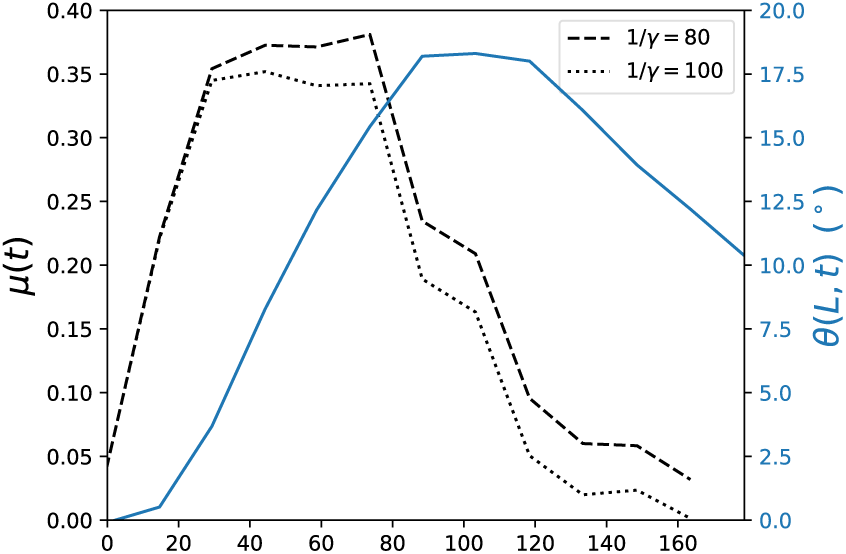
Time course of the tip angle *θ*(*L,t*) of *Arabidopsis Thaliana* seedlings as a response to a pulse of blue light [26] (solid blue line, right y-axis), as also displayed in Fig. 2a. We numerically calculate the derivative 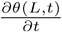, and substitute this, together with an estimated *γ*, in Eq. 4, yielding the functional form of the estimated response function, plotted against the left y-axis. The black dashed and dotted lines represents *μ*(*t*) calculated with the upper and lower bound of *γ* respectively, as detailed in the main text.

Experimental observations of plant tropisms over a century ago showing that growing plants respond by changing their shape identically to different combinations of stimuli - intermittent in time (termed summation experiments) or with reciprocal ratios of intensity and duration (reciprocity). This suggests a tropic response that is non-local in space-time. Here, we have provided a simple but general quantitative framework for the ability of growing shoots to integrate stimuli in space and time. Our theory takes the form of an integro-differential dynamical law for the growth response in terms of a memory kernel *μ*(t) and helps explain both the summation and reciprocity experiments. Using the observed response to a pulse stimulus allows us to relate the form of the memory kernel *μ*(t) to experimentally tractable variables including the temporal evolution of the angle at the tip *θ*(*L, t*) and provide upper and lower bounds for the functional form of the response function. We note that the two estimates do not deviate significantly, suggesting the our result is robust.

Our theory is but the first step in understanding how growing systems respond to environmental stimuli. Natural questions that arise include testing the assumption of linear response experimentally, comparing the response functions for phototropic and gravitropic responses, which share many signal transduction processes, and incorporating additional effects such as internal cues [28], and the role of passive drooping of a growing shoot [29], all problems for the future.

## Acknowledgments

We would like to thank Bruno Moulia and Hugo Chauvet for their help and fruitful conversations.

